# Vitrification and rapid rewarming of precision-cut liver slices for pharmacological and biomedical research

**DOI:** 10.1101/2024.12.08.627273

**Authors:** Srivasupradha Ramesh, Joseph Sushil Rao, Bat-Erdene Namsrai, Benjamin Fisher, Diane K. Tobolt, Michael Megaly, Michael L. Etheridge, Timothy L. Pruett, Davis Seelig, Paari Murugan, Bashar Aldaraiseh, Erik B. Finger, John C. Bischof

**Affiliations:** Department of Mechanical Engineering, University of Minnesota, MN, USA; Department of Surgery, University of Minnesota, MN, USA; Schulze Diabetes Institute, University of Minnesota, MN, USA; Department of Veterinary Clinical Sciences, University of Minnesota, MN, USA; Department of Laboratory Medicine and Pathology, University of Minnesota, MN, USA; Institute for Engineering in Medicine, University of Minnesota, MN, USA

**Keywords:** Vitrification, Liver slice, Cryopreservation

## Abstract

**Background and Aims:** High-throughput in vitro pharmacological toxicity testing is essential for drug discovery. Precision-cut liver slices (PCLS) provide a robust system for screening that is more representative of the complex 3D structure of the whole liver than isolated hepatocytes. However, PCLS are not available as off-the-shelf products, significantly limiting their translational potential. Cryopreservation could solve this bottleneck by effectively preserving PCLS indefinitely until their time of use. Conventional cryopreservation (slow cooling in DMSO-forming ice) results in poor PCLS viability and function and, therefore, has proven unsuitable. Here, we explore an “ice-free” cryopreservation approach called vitrification and focus on culturing and assessing PCLS for 3 days post-vitrification and rewarming, given that most acute drug toxicity tests are conducted over 24h.

**Methods:** Rat liver slices were diffusively loaded with a cryoprotective agent (CPA) cocktail consisting of EG and Sucrose. The CPA-loaded PCLS were placed on a polymer cryomesh, vitrified in liquid nitrogen (LN2), and rapidly rewarmed in CPA. The vitrified and rewarmed PCLS were subsequently cultured in a controlled volume of serum-free, chemically defined media for 3 days.

**Results:** The cryopreserved PCLS maintained high viability, morphology, function, enzymatic activity, and drug toxicity response. Results show that the vitrified PCLS perform comparably to untreated controls and significantly outperform conventionally cryopreserved PCLS in all assessments (*p* < 0.05).

**Conclusions:** Rapid vitrification and rewarming of PCLS using cryomesh enabled successful preservation and culture. This approach maintained high viability, function, enzymatic activity, and drug response for 3 days in culture, similar to controls.

**Impact and Implications:** The implications of using vitrification to store PCLS are extensive. This technology provides an exciting opportunity for the development of an “off-the-shelf” cold supply chain of human PCLS from organs declined for transplant, which are sliced, cryopreserved, and stored in a repository and available for on-demand shipping for industrial and academic biomedical research. This would also allow PCLS to become a scalable, reproducible, wide-ranging, and population-representative source of tissue that can accurately mimic in-vivo conditions of the human liver. These transformative technologies could revolutionize our practice in studying not just the metabolism of drugs but also increase our capacity to study the zonal progression of many liver diseases and conduct other exciting biomedical research.

## Introduction

The costs of bringing a new pharmaceutical to market ranges between $2.6B and $6.7B, when including capital and failure costs^1^. The decrease in successful R&D outcomes within the biopharma industry poses challenges to the global healthcare ecosystem by increasing price pressure and extending timeframes to deliver life-saving treatments to patients.

Current in-vitro drug testing and discovery models are mostly animal-derived and cell-based, and they fail to accurately predict the toxicity and efficacy of drugs in humans, leading to failures during clinical trials^2^. In vitro drug metabolic studies use less complex testbeds, which do not adequately capture the complex dynamics of the multicellular mechanisms of the human body^3,4^. This pushes us toward an R&D crossroads where the need for accurate and efficient testbeds for biomedical research is critical.

Hepatotoxicity is the most common cause of pharmaceutical failure and withdrawal from the market. Ideally, drug metabolism is studied in vivo, but this becomes prohibitive in terms of cost, accessibility of samples, and ethical concerns^5^. Alternatively, PCLS can act as a well-characterized model. PCLS maintain in-vivo architecture, cell populations, and function of the whole organ. Since their introduction 30 years ago, PCLS studies have demonstrated the ability to mimic drug pathways in humans for drug efficacy and toxicity studies^6,7^. PCLS can also be created from diseased livers, making them excellent in-vitro disease models for a wide range of biomedical research. Arguably, the biggest barrier to PCLS use is the lack of standardized protocols for preserving PCLS like those that exist for cells^6^. Improving protocols for preserving, storing, and shipping PCLS would broaden access for a variety of applications. However, like organs, the viability of PCLS rapidly degrades within hours in cold storage. Under culture, PCLS can last for only a few days, but viability and function decline with time in culture^8^.

Cryopreservation, or the storage of biosystems at ultralow temperatures, could enable the on-demand supply of PCLS for pharmacotoxicology and other biomedical applications. However, conventional freezing cryopreservation protocols using slow cooling in dilute concentrations of DMSO have proven unreliable for PCLS. While slow freezing with ice is commonly employed in cell suspensions^9^, it fails in complex tissues where ice formation disrupts cellular and tissue structures, severely compromising viability and function^10^. Indeed, the best PCLS freezing protocols published have resulted in only 70% viability after 4 hours in culture^10^.

An alternative to slow freezing is vitrification, which uses high concentrations of CPAs and high cooling rates to form a glass-like “ice-free” state. We posit that vitrification would allow for the storage of PCLS at cryogenic temperatures with minimal to no loss in their viability, function, and enzymatic activity. To successfully vitrify, we must cool the PCLS faster than the CCR (critical cooling rate required to avoid ice formation) and rewarm them faster than the CWR (critical rewarming rate required to avoid ice formation). Cryopreservation of PCLS by vitrification requires a delicate balance of having sufficient concentration of CPA to avoid ice formation under the achieved cooling and rewarming rates, while minimizing toxicity from the CPA.

In this study, we used rat liver slices as a representative biological model. We chose a CPA based on previous literature^11,12^ and performed systematic studies to optimize the CPA loading and unloading conditions. To support rapid cooling and rewarming, we utilized a cryomesh system that we previously used to successfully cryopreserve islets^13^ and other organisms^14^. Finally, we assessed PCLS viability, metabolism, function, enzymatic activity, and drug response over three days in culture.

## Material and Methods

### Preparation of PCLS

Livers were recovered from 2.5-3-month-old Sprague-Dawley rats (Charles River Labs, Wilmington, MA) and sliced into 5mm wide and 250 µm thick slices as described in the supplemental information. The University of Minnesota IACUC approved the study.

### Culturing PCLS using mesh in 6 well plate

The slices were cultured in 6-well tissue culture plates with 1 ml of media per slice per well on nylon meshes with 0.3 mm filament thickness and 500 µm pore size under normoxic conditions in a 37°C incubator. We maintained the media height above the tissue at 0.45 mm. The media composition is presented in supplemental information.

### Vitrification and rewarming of PCLS

PCLS were transferred to ETFE cryomeshes with a monofilament diameter of 25 µm and 500 µm pore size. The PCLS were loaded with the CPA in steps, as indicated in Fig 2A. Once loaded with CPA, the PCLS were placed on the cryomesh and convectively vitrified by vertical immersion in liquid nitrogen to reduce Leidenfrost boiling as previously described^14^. The PCLS were then convectively rewarmed by immersion in rewarming solution at room temperature, followed by unloading the CPA from the slices at 4°C in steps as shown in Fig.2A.

**Fig. 1.**
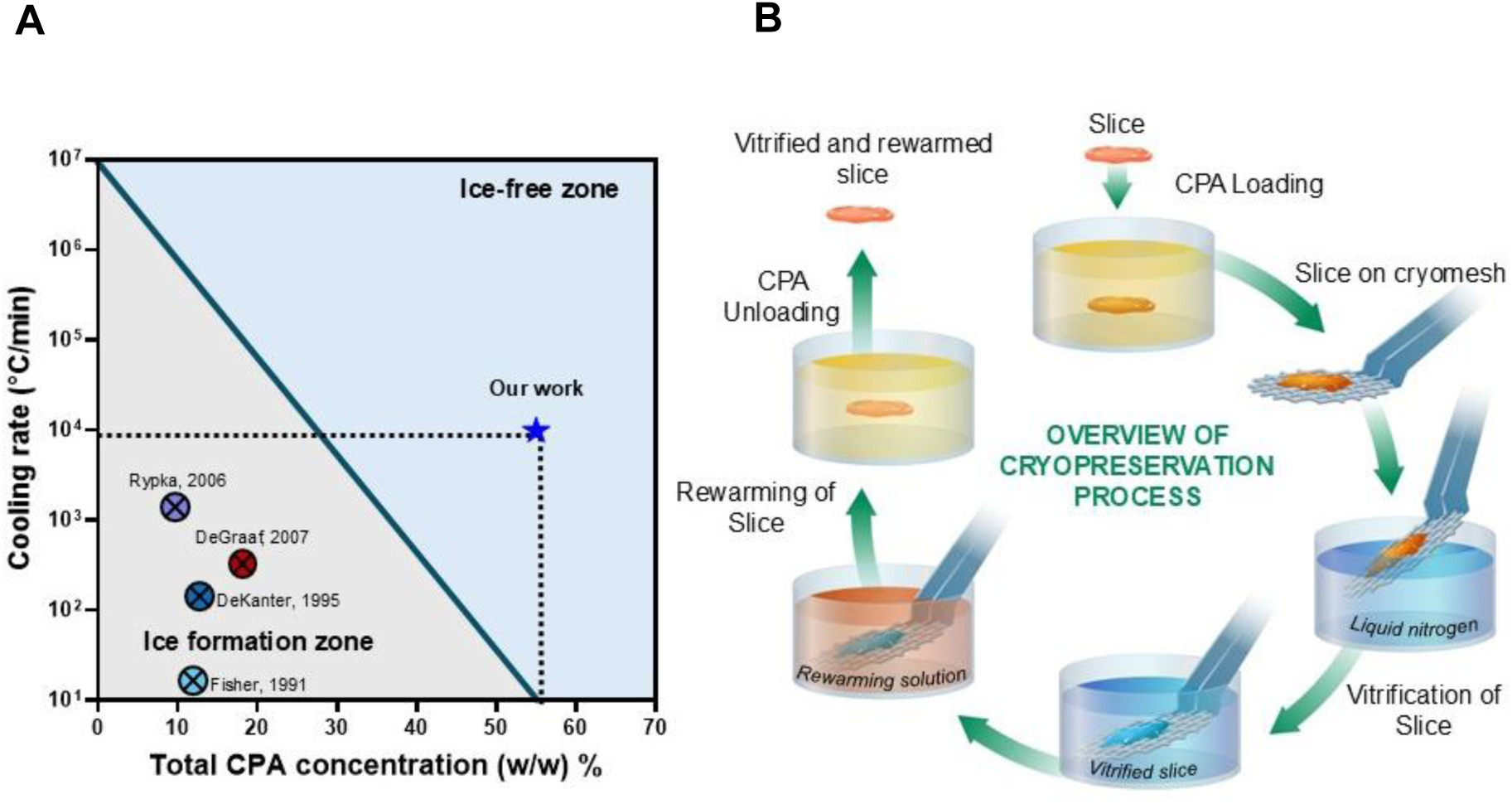
Overview of PCLS cryopreservation. (A) Previous works in the cryopreservation of PCLS report cooling rates that would lead to damaging ice formation. Our work uses high concentration cryoprotective agents and rapid cooling and rewarming to achieve ice-free vitrification. (B) Schematic diagram of the steps in the process for vitrification and rewarming using the cryomesh. CPA, cryopreservation agent.

**Fig. 2.**
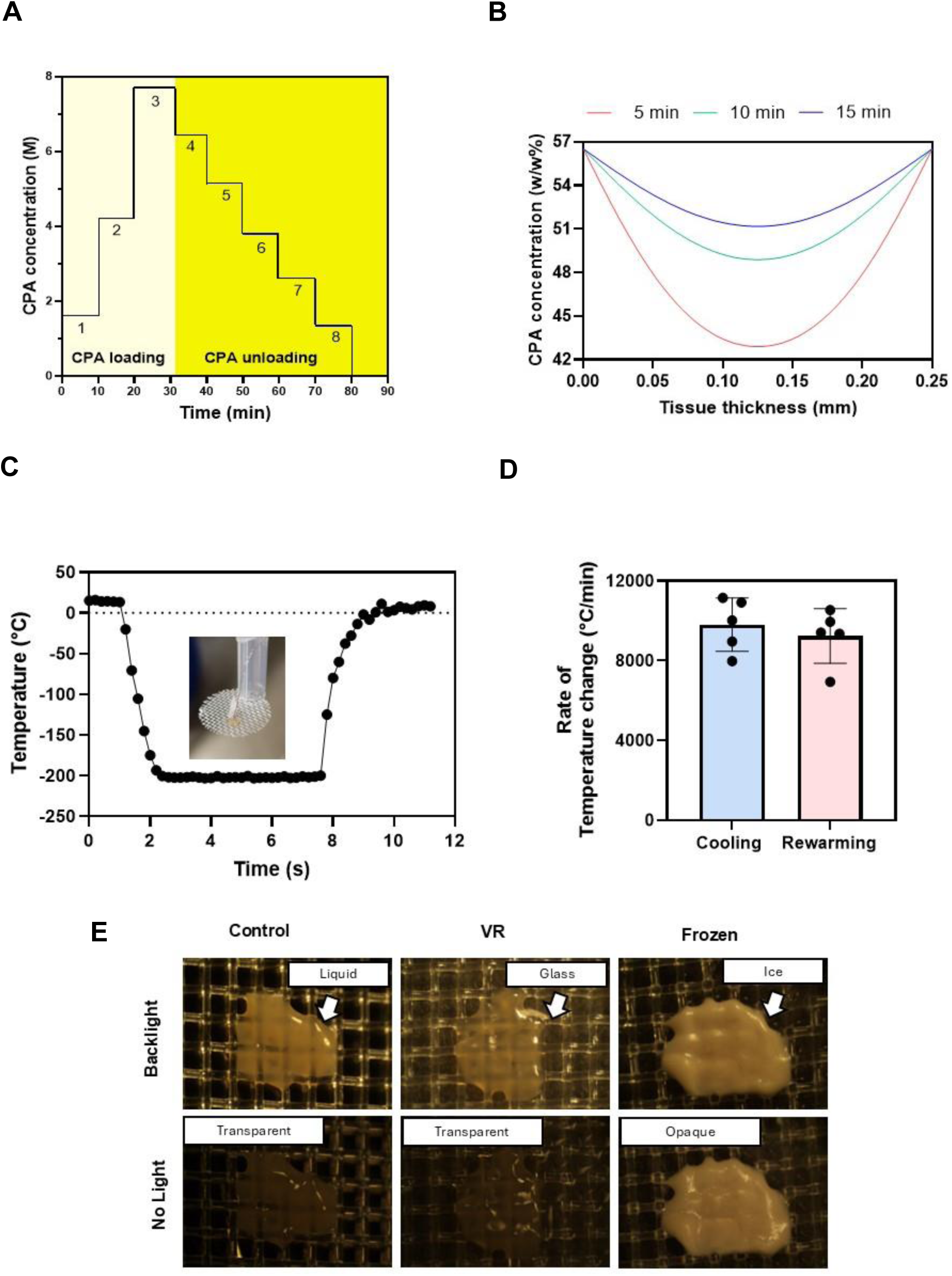
Vitrification characteristics of PCLS. (A) The CPA loading and unloading profile was an initial three steps of EG followed by a final step of 40% (v/v) EG and 0.6M Sucrose. Post vitrification, we unloaded the CPA in six steps of decreasing sucrose concentration and a final carrier solution-only step. (B) The COMSOL model estimated the total CPA concentration throughout the tissue following loading steps of 5, 10, and 15 minutes. (C) Representative thermometry of the PCLS taken during vitrification and rewarming on the cryomesh. Inset is a picture of the thermometry setup. (D) Cooling and rewarming rates measured from thermometry. (E) Visualization of ice in the PCLS. Control and VR are transparent, indicating no ice, and the frozen slice is opaque due to ice formation. CPA, cryoprotective agent; VR, vitrification and rewarming.

### COMSOL model

A Transport of Concentrated Species COMSOL (COMSOL, Burlington, MA) model was used to estimate the diffusion of CPA components for loading times. A Mixture Average diffusion model was used, and the Maxwell-Stefan diffusivities for the CPA components were calculated using the formula reported by Yu et al^15^. The coefficient for Sucrose was calculated using the viscosity values provided by the National Bureau of Standards (Circular 440)^16^ at 75 (%w/w) at 5°C. Additional details are in the supplemental information.

### Cytochrome P450 1A1 (CYP1A1) live tissue imaging and quantification

For live imaging of CYP1A1 activity, the CYP1A1 was induced in the slices using 25µM β-naphthaflavone for 24h. PCLS were then incubated with 20 µM 7-Ethoxyresorufin and 25 µM Dicumarol for 10 minutes, followed by imaging or quantification. The CYP1A1 cleaves the 7-Ethoxyresorufin to fluorescent resorufin that can be imaged and quantified^17^. The slices were imaged using an excitation wavelength of 561 nm laser in a Nikon A1RMP+ microscope. For quantification, the slices were placed in a microplate reader and imaged for 30 minutes in kinetic mode using 535/595 nm filters at 37°C. Additional details are provided in the supplemental information.

### Acetaminophen drug study with cryopreserved slices

Acetaminophen was prepared in culture media with supplements (details in supplemental information) in increasing concentrations: 0, 1, 5, 10, 20, and 50 mM. Slices were then exposed to the APAP culture media for 24 hours, after which the media was replaced with media containing no APAP and cultured for 2 more days.

### Statistics

Statistical analysis was performed in R version 4.3.2 (R Foundation for Statistical Computing, Vienna, Austria). Normality was established using the Shapiro-Wilk test to compare continuous variables, and homogeneity of variance was assessed using Levene’s test. For normally distributed group comparisons, ANOVA testing with pairwise post hoc t-test for single comparisons or Tukey HSD test for multiple comparisons were used. Non-normal variables were tested using the non-parametric Kruskal-Wallis and pairwise Wilcox (Mann Whitney U) or Dunn’s tests for individual group comparison. Grubb’s test was used to screen and censor outlier data. The Benjamini–Hochberg method was used to adjust for multiple comparisons. A *p*-value less than 0.05 was taken to be statistically significant (*) (*p* < 0.005 represented by **, *p* <0.0005 by ***, *p* < 0.00005 by ****). Continuous data are presented as mean ± standard deviation. Only statistically significant differences are shown in the figures.

### Additional materials and methods

Full details of the materials, methods, computational modeling, culture conditions, and other experimental procedures are available in the supplemental information.

## Results

### Vitrification and rewarming of PCLS

This study tested cryopreservation and post-rewarming viability, function, enzymatic activity, and drug response in 250 µm thick x 5 mm diameter PCLS from Sprague Dawley rats as a model system. We vitrified the liver slices using a cryoprotectant cocktail consisting of 40%(v/v) of EG and 0.6M Sucrose, initially developed for the cryopreservation of hepatocytes^12^, which we have used in whole rat livers^11^. A modified version of University of Wisconsin (UW) solution as carrier (starch-free with a 1 g/L concentration of PEG35k for oncotic support, as previously reported^18^).

Osmotic injury and toxicity are two of the main challenges to resolve when designing CPA loading and unloading protocols. For this study, CPA loading and unloading were performed stepwise to limit osmotic injury (Fig. 2). CPA exposure was conducted at 4°C to reduce toxicity, as we previously used in rat livers^11^.

We determined the CPA loading protocol by developing a multicomponent diffusion model in COMSOL that estimated EG and Sucrose tissue concentration over time, as previously described for other permeant CPAs^15^. The model tested three step durations (5, 10, and 15 minutes). At 5 minutes, the model predicted a minimum CPA concentration of 42.8 (% w/w) in the interior of the PCLS, whereas this increased to 48.8 (% w/w) at 10 minutes. Our previous studies^19^ estimated that a minimum of 48 (% w/w) of CPA would be needed to successfully vitrify and rewarm without ice formation at the cooling and rewarming rates achievable by our cryomesh. Thus, a 10-minute step duration was chosen for this specific CPA loading and unloading process.

The vitrification and rewarming procedure were then experimentally validated by thermometry and visual inspection. Vitrification of the slices with 10-minute loading steps indicated no visible ice (Fig. 2).

The vitrification and rewarming of the PCLS were done on ETFE cryomeshes. Temperature measurements taken during cooling and rewarming of the slices indicated average cooling rates of 9,800°C/min and rewarming rates of 9,200°C/min, which exceeded the expected CCR (24.6°C/min) and CWR (7703°C/min) at the minimum required concentration (48 w/w%) calculated from previous studies^19^.

### Viability and morphology of cryopreserved PCLS

The viability of the slices was assessed using AO/PI viability stain (live cells are green in color, and dead cells are red). Fresh control groups were compared to PCLS treated with CPA-only (CPA loaded and unloaded), vitrification and rewarming (VR), conventional cryopreservation (slow freezing and rapid thawing in cryovial after loading with 18% DMSO (v/v) for 30 minutes on ice, FT), and a negative control group (freeze-thawed 3 times with no CPA, Dead) (Fig. 3A). Live and dead cell fractions were quantified by image analysis (Fig. 3C).

**Fig. 3.**
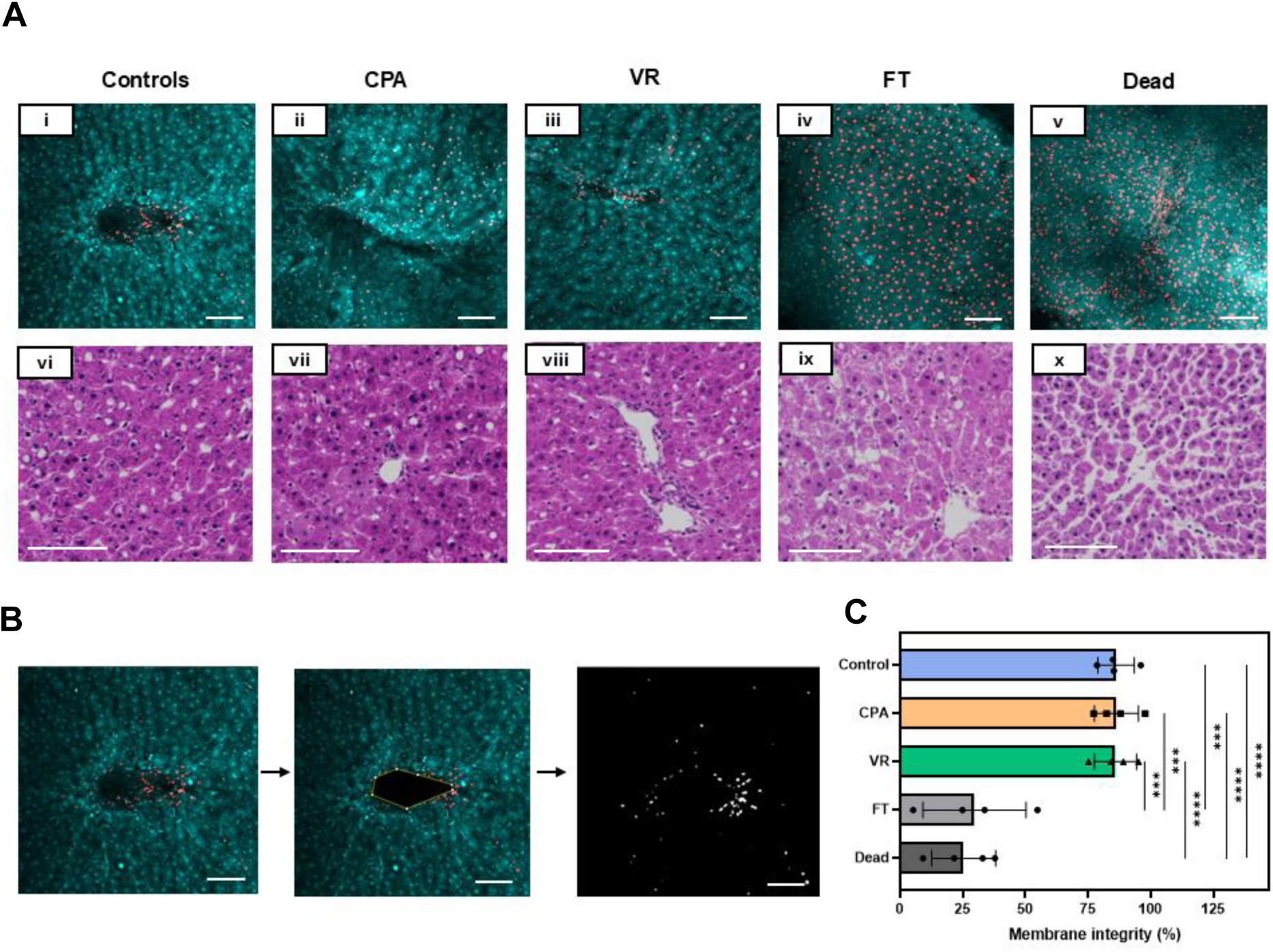
Viability and tissue morphology directly post cryopreservation. (A) (i-v) AO/PI staining where AO-stained cells appear green, and the PI-stained cells are red, indicating cell membrane compromise. (vi-x) H&E staining of the slice groups just post rewarming. (B) The viability through membrane integrity was assessed by analyzing the number of dead cells in the given z-plane by assessing areas of individual cells and comparing the number of PI quenched cells to the total number of cells. (C) Viability measure from AO/PI for each group. Levels of significance: ***, *p =0.0001;* ****, *p* <0.0001 (One-way ANOVA). AO, acridine orange; CPA, cryoprotective agent; FT, slow freezing and rapid thawing; H&E, hematoxylin and eosin; PI, propidium iodide; VR, vitrification and rewarming.

The morphology of the slices was assessed by histologic staining (H&E), which showed that PCLS from the Control, CPA, and VR groups had well-preserved hepatocytes, microarchitecture, and portal tracts. In comparison, the FT group exhibited significant vacuolization, and the Dead group showed significantly damaged microarchitecture. These were predicted to substantially affect viability, consistent with the measures observed.

### Culture conditions for liver slices

Previous attempts have been made to culture liver slices for extended periods, but for most drug discovery and testing applications, immediate acute toxicity tests are conducted within 24 hours^20^. When conducting trials for daily dose drugs, there are perspectives on how an optimal drug design for oral drugs should aim to have a half-life of 12-48h^21^. So, we set culturing timeframes to be 3 days for the initial development of the study as a conservative estimate for the practical window needed for drug studies. Previous studies have shown that sufficient passive oxygenation for hepatocytes can be provided by adjusting the media height above the cells, eliminating the need for high levels of external oxygenation^22^. Therefore, we systematically studied the height of the media above the PCLS, striving to avoid supplemental external oxygenation of the media whilst maintaining maximum viability and function. We found that 1 ml of media was required to sustain a single slice over 24 hours (200:1 media to tissue volume) and optimized the media height above the tissue to 0.45 mm. Additionally, culturing the slice on meshes enhanced oxygen diffusion from both sides. This resulted in our ability to culture PCLS with high viability, function, and drug response for up to 3 days in culture.

### Metabolic health and functional assessments in cryopreserved slices

We assessed the metabolic health of the PCLS by their ATP levels (Fig. 3A). The ATP levels of the controls, CPA, and VR groups remained high and consistent throughout the 3 days in culture (Fig. 4A). The FT group showed significantly lower ATP levels throughout the 3-day culture period, while the Dead group remained below the detection limit of the assay used (< 25nM).

**Fig 4.**
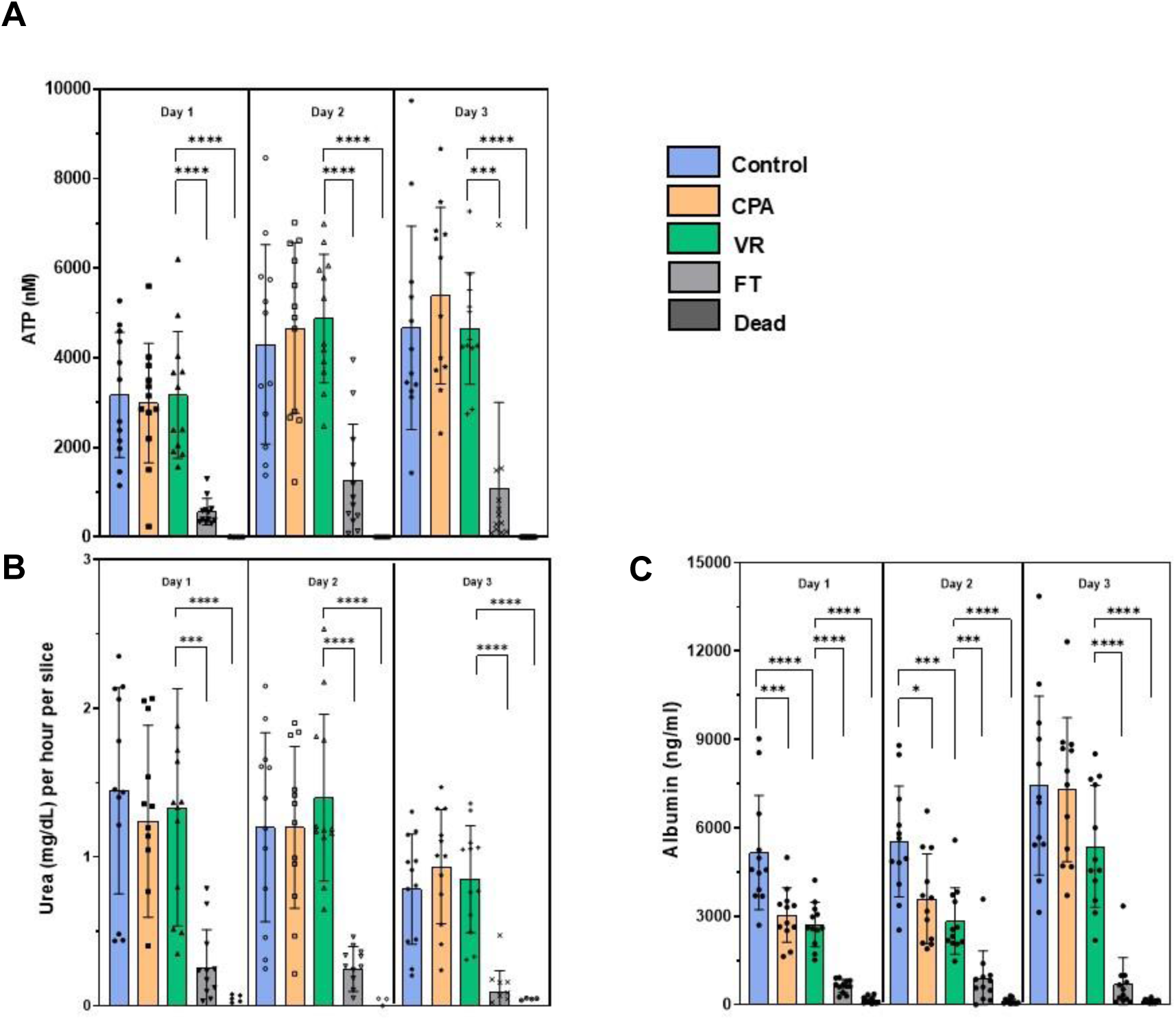
Metabolic and functional assessments of liver slices over 3 days in culture. (A) ATP assessment. (B) Urea production. (c) Albumin synthesis. Levels of significance: \**p <0.05,* ** *p* < 0.005, *** *p* <0.0005 and \*\*\*\**p* < 0.00005 (Mann-Whitney U test). CPA, cryoprotective agent; FT, slow freezing and rapid thawing; VR, vitrification and rewarming.

**Fig. 5.**
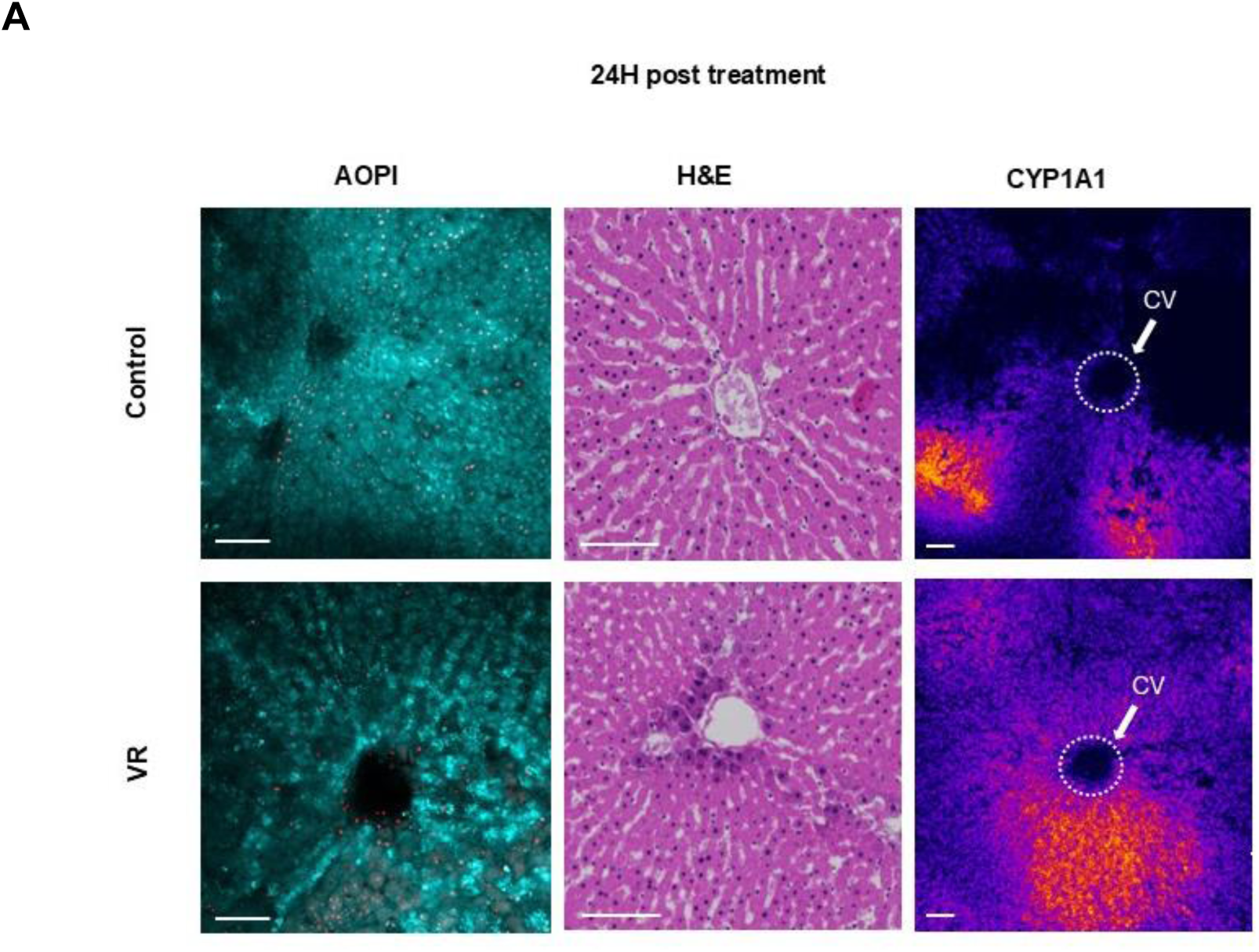
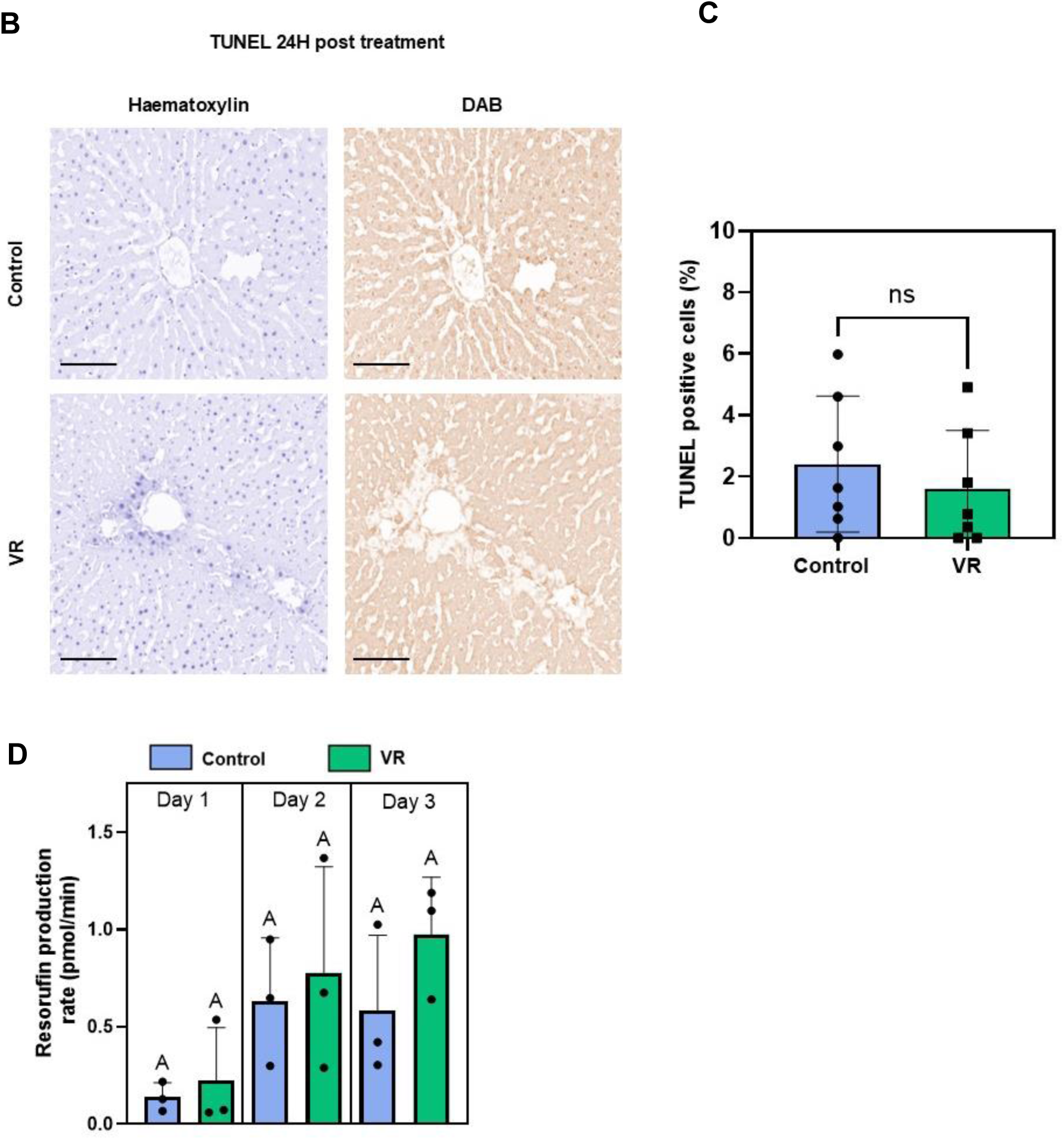
Acute injury response of slices and quantified enzymatic activity. (A) Acute evaluation of PCLS with AO/PI, H&E staining, and CYP1A1 zonated activity. (B) TUNEL staining after 24h culture to evaluate apoptosis (or necrosis) in both groups. Hematoxylin staining (blue) shows all nuclei and DAB stain (brown) shows TUNEL positive nuclei. (C) Quantification of TUNEL-positive cells (n=7). (D) Rates of resorufin production (N=3, n=4). The level of significance is presented by compact letter display (Kruskal-Walli’s test). AO, acridine orange; CYP1A1, Cytochrome P450 1A1; H&E, hematoxylin and eosin; PI, propidium iodide; VR, vitrification and rewarming; CV, central vein.

Good function of the cryopreserved slices is required for effective biomedical and drug testing applications. To measure liver slice function, we analyzed urea production (Fig. 3B) and albumin synthesis (Fig. 3C). With urea production, the CPA-only and VR groups behaved similarly to Controls, where urea production remained consistent throughout the 3 days in culture with mean values equal to or above 0.7 mg/dL for all three groups. A slight drop in urea production was observed on day 3 but was not statistically significant. The FT group showed significantly less urea production over all 3 days, consistent with other measurements. The dead group showed no measurable urea production.

Albumin synthesis stayed high and consistent with the control groups over all three days (Fig. 3C). The CPA and VR groups experienced an initial decrease in albumin synthesis compared to controls but then caught up to controls on the third day of culture. This could be due to recovery of the slices from an initial stress response on exposure to the CPA and during vitrification/rewarming, but it did not seem to hinder the ability of the slices to synthesize new albumin^23^. An increase in albumin synthesis was seen with the controls, CPA, and VR groups when comparing day 1 to day 3, indicating preserved function of the CPA and VR slices. The FT and dead groups had significantly less albumin synthesis over all 3 days in culture. However, the dead controls still showed low albumin levels, possibly indicating a slow leakage of pre-formed albumin rather than actual synthesis.

### Acute testing for zonated CYP activity, viability and apoptosis

Liver slices afford the opportunity to assess the zonal activity of cytochrome P-450 (CYP) enzymes that form the crux of xenobiotic metabolism. These enzymes are zonated, with the most activity present in zone 3 near the central hepatic vein draining the hepatic lobule. To understand if the zonal CYP activity is observed in the VR slices, we induced CYP1A1 in the control and VR PCLS for 24h, followed by imaging. This allowed for assessment of zonal CYP activity after cryopreservation where the tissue may undergo significant change and injury. We also quantified the activity throughout the 3 days in culture.

By confocal imaging, the cleaved resorufin was visible as a bright signal near the central vein in the acini of the liver slices in both the Control and VR groups, indicating zonated CYP activity in zone 3. Quantification of fluorescence demonstrated that resorufin production rate was induced and similar between control and VR PCLS over the 3-day culture. Both assessments showed that CYP activity post cryopreservation was zonated in the acute 24-hour period and that activity was maintained during all 3 days in culture (Fig 2A).

TUNEL staining was also performed after 24h to assess induction of apoptosis (or necrosis). The TUNEL-positive cell fraction did not differ between VR and control PCLS (Fig. 3BC). Control and VR groups showed an average of 2.8% and 1.8% TUNEL-positive cells, respectively, which was not statistically different.

### Acetaminophen (APAP) hepatotoxicity in fresh and cryopreserved PCLS

The most critical issue related to in vitro drug studies is the potential for hepatotoxicity, representing a major failure mechanism for pharmaceuticals. To demonstrate the use of cryopreserved PCLS as an in-vitro model for drug testing, we exposed the control, VR, and Dead slices to different concentrations of APAP. We chose APAP as a model drug for testing since it is well-studied, and APAP overdose is a leading cause of hepatotoxicity, which in some cases could lead to fatal acute liver failure^24^.

Liver slices were exposed to increasing concentrations of APAP for 24 hours. The APAP dosage was chosen based on previous studies in liver cell lines ^25^. Urea was measured in the same slices over 3 days in culture (Fig. 6B). The slices were homogenized at the end of 72 hours to measure ATP content (Fig. 6A). VR and Control slices performed very similarly, maintaining function and viability APAP concentrations to 5 mM, but 10 mM APAP resulted in severe injury and loss of function that increased over the culture period. At 20 mM and 50 mM APAP exposure, complete necrosis of slices occurred even on day 1, with no urea detectable. The ATP content in the slices at the end of day 3 also showed similar trends, with the slices exposed to 0-5 mM APAP maintaining ATP content while the slices exposed to 10-50 mM APAP showed undetectable ATP levels (below 25 nM).

**Fig. 6.**
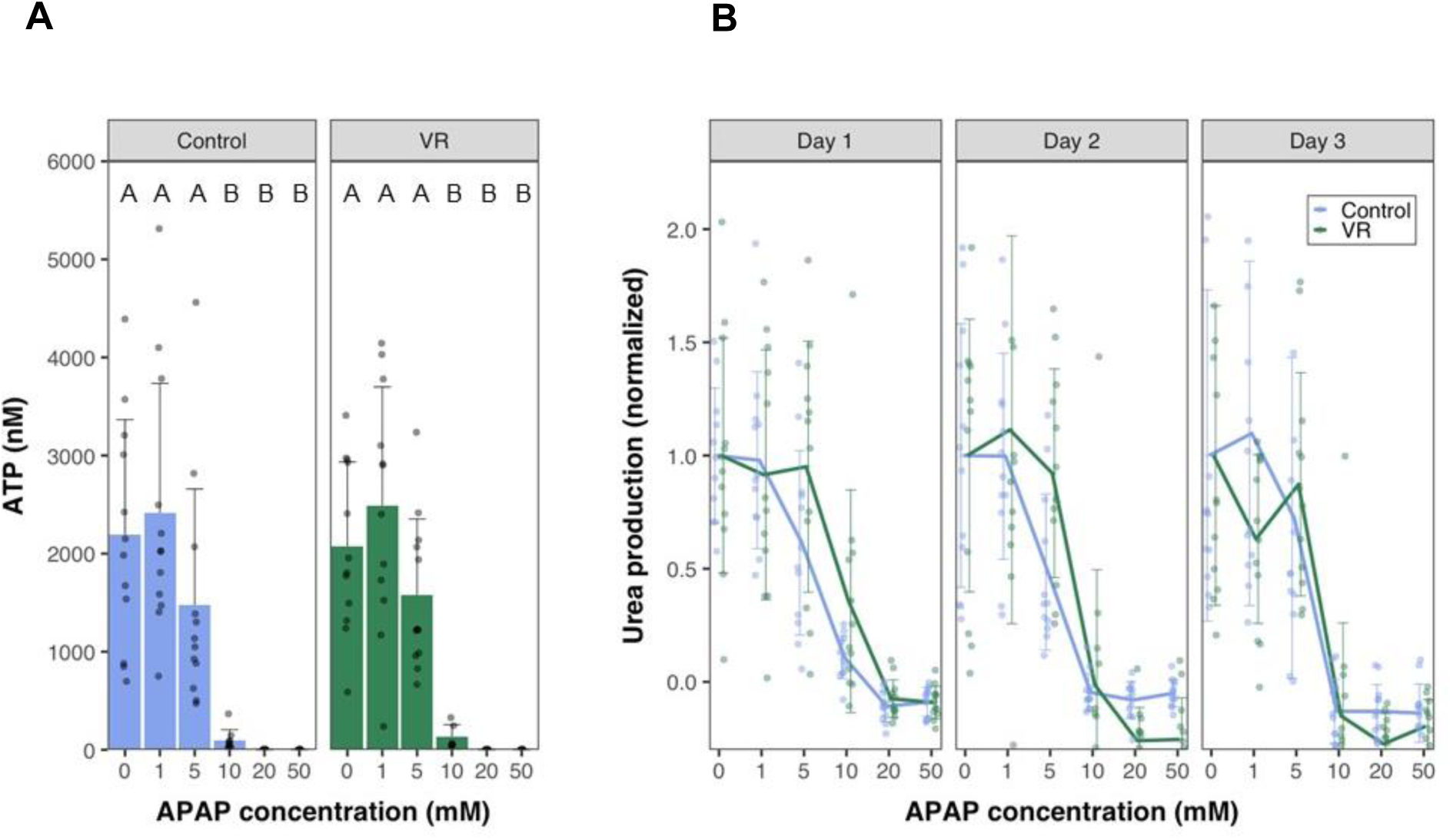
Drug response of PCLS. (A) ATP levels at the end of day 3 in culture of control and VR slices exposed to varying concentrations of Acetaminophen (APAP). (B) Urea levels measured normalized to 0mM concentration spanning a 3-day culture period on exposure to different APAP concentrations. Data are mean ± standard deviation. Levels of significance are represented by compact letter display (Dunn’s test). APAP, N-acetyl-para-aminophenol (acetaminophen), VR, vitrification, and rewarming.

## Discussion

Preventing drug failures during human clinical trials requires careful evaluation during the pre-clinical stage. This capability can prevent patient harm, eliminate ineffective therapies, and ultimately increase successful outcomes while lowering costs. In the absence of in vivo testing, which is cost-prohibitive and not fully accessible, cryopreserved liver tissue slices are a promising solution. PCLS banking from representative patient populations would open the possibility for repeatable drug screening and provide an accurate and patient-representative model for studying diseases and their progressions. In this study, we illustrated in detail the first successful cryopreservation of rat liver slices through ice-free vitrification and rewarming to maintain viability, function, and enzymatic activity and show drug response post-rewarming out to 3 days.

As mentioned, we focused on the culture of liver slices for 3 days, an acute testing window for a wide range of biomedical applications. Notably, no current standardized methodology for the culturing of liver slices exists. Many culturing methods involve using large Erlenmeyer flasks and culturing the slices in external oxygenated environments using carboxygen^26^. Even with continuous improvement in culturing technology, the viability with most PCLS culture has only been reported for 24-48h^27,28^. The most extended culture of PCLS were reported by Parish et al., who cultured PCLS for 6 days using a bioreactor^8^, and Wu et al., who reported culturing PCLS for 15 days^29^. However, they reported a decrease in specific gene expressions over the days in culture, which may indicate a transition to a non-representative sample. Inspired by air-interface culturing systems, we achieved a reproducible culture of liver slices for up to 3 days using the simple technique of suspending the PCLS on nylon meshes—this and optimizing the media height above the tissues allowed for better oxygen diffusion into the tissue.

Having established a culture approach that can be reliably replicated, we turned to cryopreservation. Unsuccessful previous studies on liver slices employed slow rates of freezing and thawing (<2,000°C/min cooling and rewarming) in cryovials using low concentrations of DMSO (18-22 %v/v)^10^. These methods resulted in poor viability and function of slices even after short-term culture. The poor viability could be attributed to disruption of the cell membranes due to ice formation and damage to the necessary extracellular microarchitecture disrupting normal function. Another major challenge with conventional methods for PCLS cryopreservation is a lack of reproducibility, given the limited control over where ice formation occurs. Vitrification circumvents these issues and allows for preserving PCLS for indefinite periods. Until now, limited attempts at vitrification and rewarming liver slices have been reported^30^, but the maintenance of viability and function of tissues over multiple days in culture has not been demonstrated.

Selecting the right CPA and protocol for loading and unloading were critical to succeed at vitrification and rewarming, as injury can manifest in the form of osmotic injury and direct CPA toxicity^31^. Osmotic injury occurs when the cells are exposed to sizeable transmembrane concentration gradients of the CPA, which causes acute shrinking of cells during loading and swelling of cells during unloading, leading to injury. One of the mechanisms of toxicity is the metabolization of the CPA components by the cells and their conversion into toxic byproducts. Osmotic injury can be minimized by controlled loading and unloading of the CPA, mitigating acute shrink-swell behavior. We reduced toxicity by lowering the temperature of CPA exposure, which slows the metabolism of the components by the cells. Hence, we loaded the PCLS stepwise at 4°C to minimize injury and vitrified and rewarmed slices at high cooling and rewarming rates using a cryomesh. The VR slices demonstrated markers comparable to fresh slice controls for all metrics assessed in this study. This observed behavior was consistent for the VR slices over 3 days in culture.

One of the critical differences that the PCLS provide as in-vitro tools over cells is their ability to capture zonal activity during xenobiotic liver function. In this study, CYP1A1 activity was similar and sustained for control and VR PCLS for all days in culture. This has important implications for drug testing protocols and understanding the progression of liver diseases. The initiation and progression of liver diseases also occur in a zone-wise manner, with many non-alcoholic and alcoholic diseases of the liver beginning in zone 3 and progressing to zone 1^32^. These zone-dependent disease and signaling pathways cannot be studied using conventional cell-based in-vitro models.

The use of VR PCLS for drug hepatoxicity was demonstrated with the common toxin, APAP, with demonstration of dose-dependent toxicity similar to control PCLS. The response of the VR group to APAP was similar to Controls, indicating no significant differences between the two groups on a macroscopic level over 3 days of culture. Most prior APAP toxicity testing outlined in the literature was performed using liver cell models. Increased toxicity levels have been reported at primarily high concentrations (>10 mM), but injury also occurs at lower concentrations (≤ 5 mM), where mitochondrial injury is thought to be the leading cause ^33–35^. In this PCLS study, there was no significant difference in ATP levels on APAP exposure from 0 mM to 5 mM in both groups. This indicates the opportunity to research the mechanisms of APAP toxicity using PCLS, which would more closely resemble in-vivo whole liver conditions. To summarize, this study used rat liver slices as a model system to establish successful vitrification and rewarming using the cryomesh. Future studies will focus on obtaining human tissue to demonstrate success with both normal and diseased human liver slices. Overall, our study offers exciting opportunities to expand the use of PCLS in pharmacology and hepatology.

## Acknowledgments

The authors would like to acknowledge Paula Overn, Colleen Forster, and Adam Lewis for performing histological staining, Robin Francis for editorial assistance, and Andy Grams for technical illustration. This work was also supported by the resources and staff at the University of Minnesota University Imaging Centers (UIC). SCR_020997.

## Author contributions

J.S.R. contributed to obtaining AO/PI imaging data, B.N. and M.M. contributed to the recovery of livers, B.F. contributed with assay data collection, D.S., P.M., and B.A. contributed to histological data interpretation, T.P. contributed to experimental design, M.E. contributed to experimental design and manuscript writing, J.C.B. and E.B.F. contributed to experimental design and manuscript writing and contributed as senior and corresponding authors.

## Abbreviations

PCLS: Precision cut liver slices
CPA: Cryopreservation agent
DMSO: Dimethyl Sulfoxide
EG: Ethylene Glycol
CCR: Critical Cooling Rate
CWR: Critical Warming Rate
ETFE: Ethylene tetrafluoride
UW: University of Wisconsin
AO/PI: Acridine Orange/Propidium Iodide
VR: Vitrified and Rewarmed
APAP: N-acetyl-para-aminophenol (acetaminophen)
DAB: 3,3′-diaminobenzidine

