## Supplementary material for "Vitrification and rapid rewarming of precision-cut liver slices for pharmacological and biomedical research": Supplemetal information

### Supplementary Information

#### Methods and Materials

##### Preparation of PCLS

A 5 mm biopsy punch was used to obtain biopsies of the liver, followed by embedding of the biopsy onto the specimen tube using 3% (w/v) of low-melting agarose (IBI Scientific, IB70051) prepared in phosphate buffered saline (PBS). The specimen tube was then cooled using a chilling block to quickly set the agarose, followed by slicing using a Compresstome (Precisionary Instruments, MA, USA). The buffer tray of the Compresstome was filled with ice-cold PBS, and cut slices were collected and placed in cold modified UW solution until further use (2-3hours).

##### Culture media composition

The culture media was WilliamsE media (Gibco) supplemented with 10,000 ng/ml Insulin, 5.5 µg/ml Transferrin, 5 ng/ml Selenium (all Sigma), 2 mM L-glutamine (Sigma), 10 mM HEPES (Gibco), 50 µg/ml Gentamicin (Sigma), 2.5 µg/ml amphotericin-B (Sigma), and 0.1 µM Dexamethasone (Sigma). No serum was added to the culture media.

##### COMSOL model

COMSOL Multiphysics 5.6 was used to model the CPA diffusion in PCLS. The preset properties for human liver were used for the model with heat capacity $C_p = 3540$ J/(kg K), density = 1079 kg/m$^3$, thermal conductivity = 0.52 W/(m K). A 0.25 mm square geometry was used to represent the tissue, as the characteristic length of interest is the thickness of the tissue. The Transport of Concentrated Species physics model was used with species diffusivity as follows: EG-Sucrose → 5.226e-12 m$^2$/s, EG-carrier → 3.85e-11 m$^2$/s and Sucrose-carrier → 5.618e-12 m$^2$/s. The temperature was set to 277 K with a mixture density of 1140 kg/m$^3$ and molar mass of EG, Sucrose and carrier to 0.062, 0.34 and 0.018 kg/mol respectively. An initial value of 4.2 M EG was set with 0 M Sucrose, and an inflow of 7.1 M EG and 0.6 M Sucrose was set as the boundary conditions on both ends of the tissue to estimate the final step time of CPA diffusion required for successful vitrification. The concentrations were then obtained for different timepoints of 5, 10, and 15 minutes. This is represented in Fig. 2.

##### Assay assessments

Assays were used for urea, albumin, and ATP as follows. For the assessment of urea, the slices were transferred to 24 well plates containing 0.25 ml of urea media for incubation times ranging from 1.5-3 hours. The urea media consisted of a KHB base solution with 10 mM ammonium chloride (Sigma Aldrich) and 2 mM L-ornithine (Sigma Aldrich) to assist in the initiation of the urea cycle. The slices were then either snap frozen, stored, and then homogenized for ATP assessment or put back into culture with fresh culture media in incubators at 37 °C with 5% $CO_2$. QuantiChrom Urea Assay Kit (BioAssay Systems) was used for the urea assay. Albumin was assessed from the culture media using an ELISA assay (Rat albumin ELISA Kit, ICL). Finally, Roche Bioluminescence Assay Kit CLS II was used for ATP assay. All assays were performed according to the manufacturer's instructions.

For AO/PI live/dead images presented in Fig.3, the PCLS were incubated with 8 ng/ml AO and 20 ng/ml PI (Millipore Sigma) for 5 min at room temperature. They were then imaged with an Olympus Fluoview 3000 inverted confocal microscope (Olympus) with 502/525-nm filters for AO and 493/636-nm filters for PI. The PCLS images were captured at 4,020 × 4,020-pixel resolution using a 20× magnification objective.

**Image analysis using ImageJ**

With ImageJ, we obtained a relative viability measure by counting the average number of dead cells (stained with red nuclei) to the total area by measuring the average area of each cell and allowing to estimate the number of cells in the given z-plane by excluding the central vein area and dividing the total area by the area of each cell. This was then followed by converting the red channel (PI signal) to 8-bit, followed by thresholding and counting the number of red nuclei using particle analysis, setting a threshold of 10 $pixel^2$ area. The membrane integrity is presented as a percent using this image analysis method in Fig.3C.

**CYP1A1 (Resorufin production) live imaging and quantification**

For live imaging of CYP1A1 activity, the slices were incubated with culture media containing 25 µM β-naphthaflavone made in DMSO with 2.1 (v/v)% final DMSO concentration) for 24 hours to induce CYP1A1. They were then incubated with WilliamE media (no phenol red), 20 µM 7-Ethoxyresorufin, and 25 µM Dicumarol for 10 minutes and imaged using an excitation wavelength of 561nm laser in a Nikon A1RMP+ microscope. The CYP1A1 cleaves the 7-Ethoxyresorufin to fluorescent resorufin that can be imaged and quantified[22]. For quantification, the slices were placed in a microplate reader (Synergy HT, BioTek) and imaged for 30 minutes in kinetic mode using 535/595 nm filters at 37°C.

**Reagents and Equipment:**

| Equipment/Reagent | Company | Location |
|---|---|---|
| Compresstome Model VZ-310-0Z | Precisionary Instruments | Ashland, Massachusetts, USA |
| β-naphthaflavone | Cayman Chemical Company | Ann Arbor, Michigan, USA |
| 7-Ethoxyresorufin | | |
| Dicumarol | | |
| Acetaminophen | | |
| ETFE Cryomesh Cat# 64700-24 | Electron Microscopy Sciences | Hatfield, Pennsylvania, USA |
| Nylon culturing mesh Cat #9318T44 | McMaster Carr | Elmhurst, Illinois, USA |
| Microplate reader Synergy HT | BioTek | Winooski, Vermont, USA |
